# An easy and reliable whole blood freezing method for flow cytometry immunophenotyping and functional analyses

**DOI:** 10.1101/2020.08.12.248450

**Authors:** Cecile Braudeau, Nina Salabert-Le Guen, Chevreuil Justine, Rimbert Marie, Jerome C. Martin, Regis Josien

**Affiliations:** Laboratoire d’Immunologie, CIMNA, LabEx IGO “Immunotherapy, Graft, Oncology”, F-44000 Nantes, France; CHU Nantes, Nantes Université, Inserm, Centre de Recherche en Transplantation et Immunologie, UMR 1064, ITUN, F-44000 Nantes, France

## Abstract

**Background:** Immune profiling by flow cytometry is not always possible on fresh blood samples due to time and/or transport constraints. Besides, the cryopreservation of peripheral blood mononuclear cells (PBMC) requires on-site specialized lab facilities, thus severely restricting the extent by which blood immune monitoring can be applied to multicenter clinical studies. These major limitations can be addressed through the development of simplified whole blood freezing methods.

**Methods:** In this report, we describe an optimized easy protocol for rapid whole blood freezing with the CryoStor^®^ CS10 solution. Using flow cytometry, we compared cellular viability and composition on cryopreserved whole blood samples to matched fresh blood, as well as fresh and frozen PBMC.

**Results:** Though partial loss of neutrophils was observed, leucocyte viability was routinely >75% and we verified the preservation of viable T cells, NK cells, monocytes, dendritic cells and eosinophils in frequencies similar to those observed in fresh samples. A moderate decrease in B cell frequencies was observed. Importantly, we validated the possibility to analyze major intracellular markers, such as FOXP3 and Helios in regulatory T cells. Finally, we demonstrated good functional preservation of CS10-cryopreserved cells through the analysis of intracellular cytokine production in ex vivo stimulated T cells (IFNg, IL-4, IL-17A,) and monocytes (IL-1b, IL-6, TNFa).

**Conclusions:** In conclusion, our protocol provides a robust method to apply reliable immune monitoring studies to cryopreserved whole blood samples, hence offering new important opportunities for the design of future multicenter clinical trials.

## INTRODUCTION

Major obstacles faced by multicenter studies to including systematic immune monitoring have been sample quality standardization and time between drawing and lab analysis. Flow cytometry analysis 4-to-24h post blood collection was shown to generate inaccurate phenotypic information because of the progressive loss of membrane marker expression (1) (2). While commercially available collection tubes with stabilizing reagents such as Cyto-Chex (Streck Cyto-Chex™, La Vista, NE) or TransFix^®^ (Cytomark, Buckingham, UK) were developed to overcome time constraints (3) (4), the expression of several membrane markers was affected by fixation, especially in myeloid cells (5). Moreover, these methods were not suitable for intracellular protein staining or functional analysis, and storage duration could not exceed a few days (6). In most multicenter studies, immune monitoring requires peripheral blood mononuclear cells (PBMC) isolation. However, even if PBMC can be cryopreserved for later analysis, polymorphonuclear cells (PMs) are lost. In addition, PBMC isolation with density gradients like Ficoll has been reported to induce biased distributions of specific immune cell subsets such as CD8+ T cells (7). Besides, PBMC preparation requires rapid access to a specialized lab for processing by well-trained staff.

A limited number of studies reported about whole blood freezing methods but, to our knowledge, none described the possibility to include intracellular staining and functional analyses. In this study, we sought to develop a simple method for rapidly freezing whole blood samples with the commercially available CryoStor^®^ CS10 freezing solution, and to validate the possibility to reliably analyze these samples by flow cytometry. By comparing to fresh whole blood as well as to fresh and frozen PBMC, we showed that our method preserved membrane marker expression and allowed for intracellular staining as well as functional assays.

## MATERIALS AND METHODS

### Biological samples

Venous blood samples (EDTA or lithium heparin tubes) were obtained from 5 healthy donors from Etablissement Français du Sang (EFS, Pays de la Loire, Nantes, France). All blood samples were processed within 3 h after collection.

### Sample preparation

Peripheral blood mononuclear cells (PBMC) were isolated by Ficoll-Paque density gradient centrifugation (GE Healthcare Europe, Velizy, France). PBMC were immediately processed for flow cytometry analysis or frozen in fetal bovine serum/7.5% DMSO (Sigma, Saint Louis, USA) and conserved in liquid nitrogen for 3 months. For whole blood freezing, ice-cooled whole blood was mixed with ice-cooled CryoStor^®^ CS10 freezing solution (Biolife Solutions, StemCell, Grenoble, France) at ratio 1:18. Cryovials were stored in an ice-cooled CoolCell freezing container (BioCision) and conserved at −80°C during 3 months. PBMC and frozen whole blood were thawed in a 37°C water bath, washed with pre-warm PBS and resuspended in PBS for immediate use.

### Immunophenotyping

Cells were stained for flow cytometry analysis using DuraClone tubes (Beckman Coulter, Roissy CDG, France) containing fluorochrome-conjugated dried antibodies (Table 1), as follows: DuraClone IM phenotyping Basic (CD45, CD3, CD4, CD8, CD14, CD16, CD19, CD56), DuraClone IM Dendritic Cell (CD45, CD1c, CD11c, CD16, CD123, HLA-DR, CLEC9A, Lineage cocktail: CD3, CD14, CD19, CD20, CD56), DuraClone IM Granulocytes (CD45, CD11b, CD15, CD16, CD33, CD62L, CD274, CD294, Lineage cocktail: CD3, CD14, CD19, CD56), DuraClone IM Treg (CD45, CD3, CD4, CD25, CD39, CD45RA, FOXP3, Helios). For the latter, an anti-CD127 antibody (BD Biosciences, Le Pont de Claix, France) was also added. Stainings were performed according to manufacturer instructions. Briefly, we used 50μl of fresh/frozen whole blood or 0.5×10^6^ PBMC for the Treg panel, 100μl of fresh/frozen whole blood or 1×10^6^ PBMC for Basic and Granulocytes panels, and 200μl of fresh/frozen whole blood or 2X10^6^ PBMC for the Dendritic cell panel. The Perfix-nc kit (Beckman Coulter) was used for Treg tube. A viability dye was included in all Duraclone panels (Live/dead fixable dead cell stain kit from ThermoFisher Scientific or DAPI (4’,6-Diamidino-2-Phenylindole, Dilactate from ThermoFisher Scientific (Courtabœuf, France)). Labeled cells were acquired on the cytometer Cytoflex (Beckman Coulter), data were analyzed with Kaluza software V2.1 (Beckman Coulter).

**Table 1:**
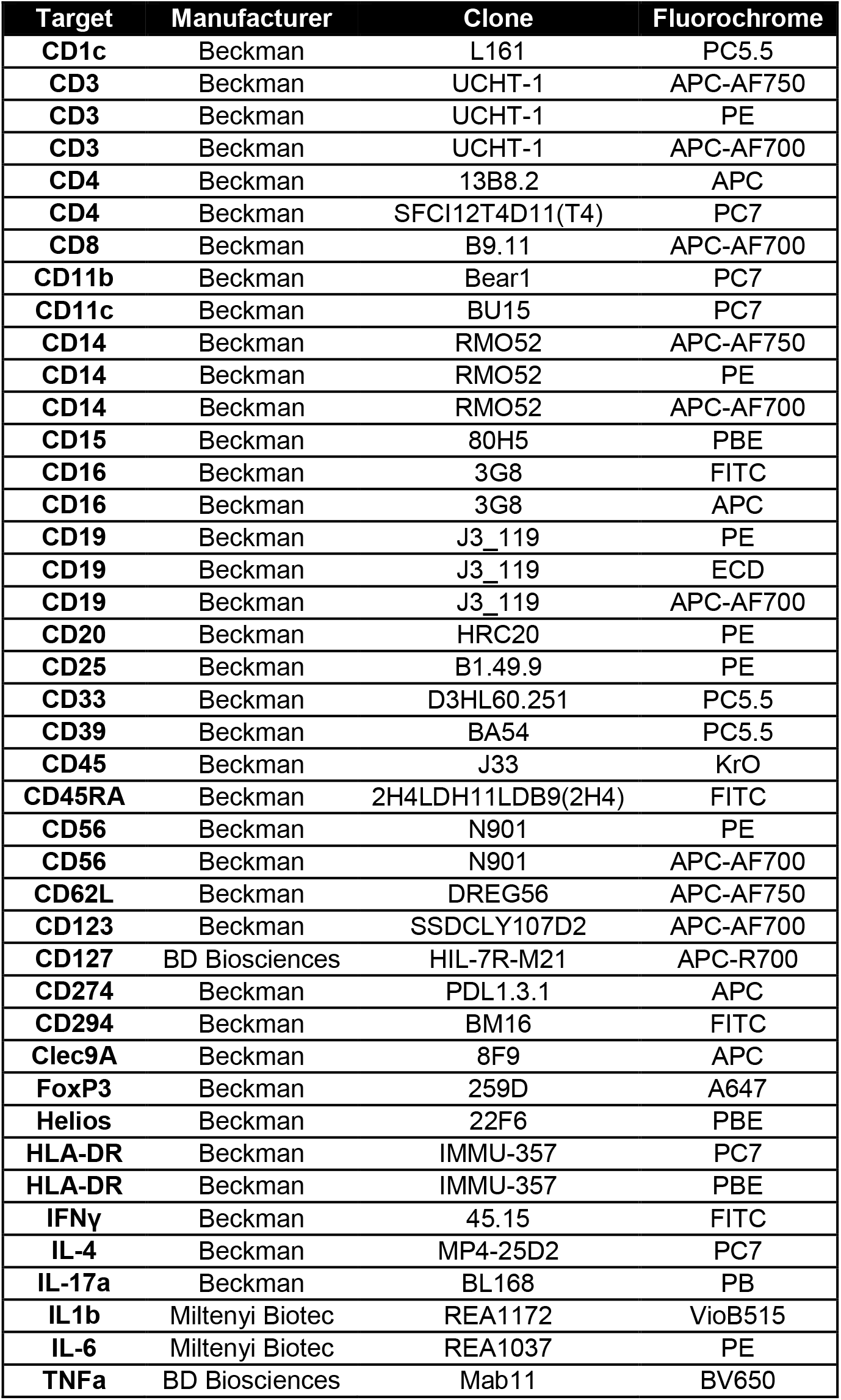
List of antibodies.

### T cell cytokine production

Intracellular cytokine production by T cells was analyzed after stimulating PBMC or whole blood (fresh or frozen) 4h at 37°C in DurActive1 tubes (Beckman Coulter), which contain PMA, ionomycin and brefeldin A. Cells were stained with a viability dye, LIVE/DEAD™ Fixable Red Stain Kit, during 30min and then transferred into a DuraClone IF T Helper cell tube (CD3, CD4, IL-4, IL-17A, IFNγ) (Beckman Coulter), using the IntraPrep fixation permeabilization kit (Beckman Coulter). Cells were further stained with an anti-CD45RO antibody (Beckman Coulter).

### Monocyte cytokine production

Intracellular cytokine production by monocytes was assessed after stimulation of fresh or frozen blood during 4h at 37°C with 1μg/ml of lipopolysaccharide (LPS) (Sigma Aldrich, St Quentin Fallavier, France). Brefeldin A (5μg/ml, Sigma Aldrich) was added during the last 3h of culture. After stimulation, cells were stained with a viability dye, LIVE/DEAD™ Fixable Red Stain Kit (ThermoFisher Scientific) and antibodies against membrane markers were added (CD45, CD14, HLA-DR from Beckman Coulter). Intracellular staining was performed using the IntraPrep fixation permeabilization kit (Beckman Coulter) and IL-1b, IL-6 mAbs from Miltenyi Biotec (Paris, France), and TNFa mAb from BD Biosciences.

### Statistical analyses

All statistical analyses were performed using GraphPad prism 5.01 software (GraphPad Software, San Diego, CA, USA). The Friedman non-parametric test with post-hoc Dunn’s multiple comparisons were used for one-way analysis of variance (ANOVA). Paired comparisons of PMs and of monocyte stimulation were done with the Wilcoxon t-test. p-values<0.05 were considered for statistical significance.

## RESULTS

### A rapid and easy-to-use whole blood freezing method

We set up an easy-to-use method for whole blood freezing, which requires minimal handling and time (<15 min), as compared to PBMC isolation (**Figure 1**). This method consists in slowly mixing freshly collected ice-cooled blood with ice-cooled CryoStor^®^ CS10 freezing medium at ratio 1:18 before transfer into cryotubes and storage in an ice-cooled CoolCell freezing container. CryoStor^®^ CS10 is a protein- and animal component-free freezing medium based on the HypoThermosol® formulation and containing 10% dimethyl sulfoxide (DMSO). This product has been initially developed for cell and tissue freezing (8,9) but to our knowledge its use for whole blood has not been described. As no particular equipment is needed, this method allows whole blood freezing at patient’s bedside and is well-suited for clinical centers without rapid access to lab facilities. Although this process requires rather large volume of freezing solution, the overall cost is balanced by the minimal time of work required before freezing, when compared to PBMC preparation. To validate this method, cryotubes were stored for 3 months at −80°C before flow cytometry analysis. All analyses were done in comparison to fresh blood as well as matched fresh and frozen PBMC.

**Figure 1:**
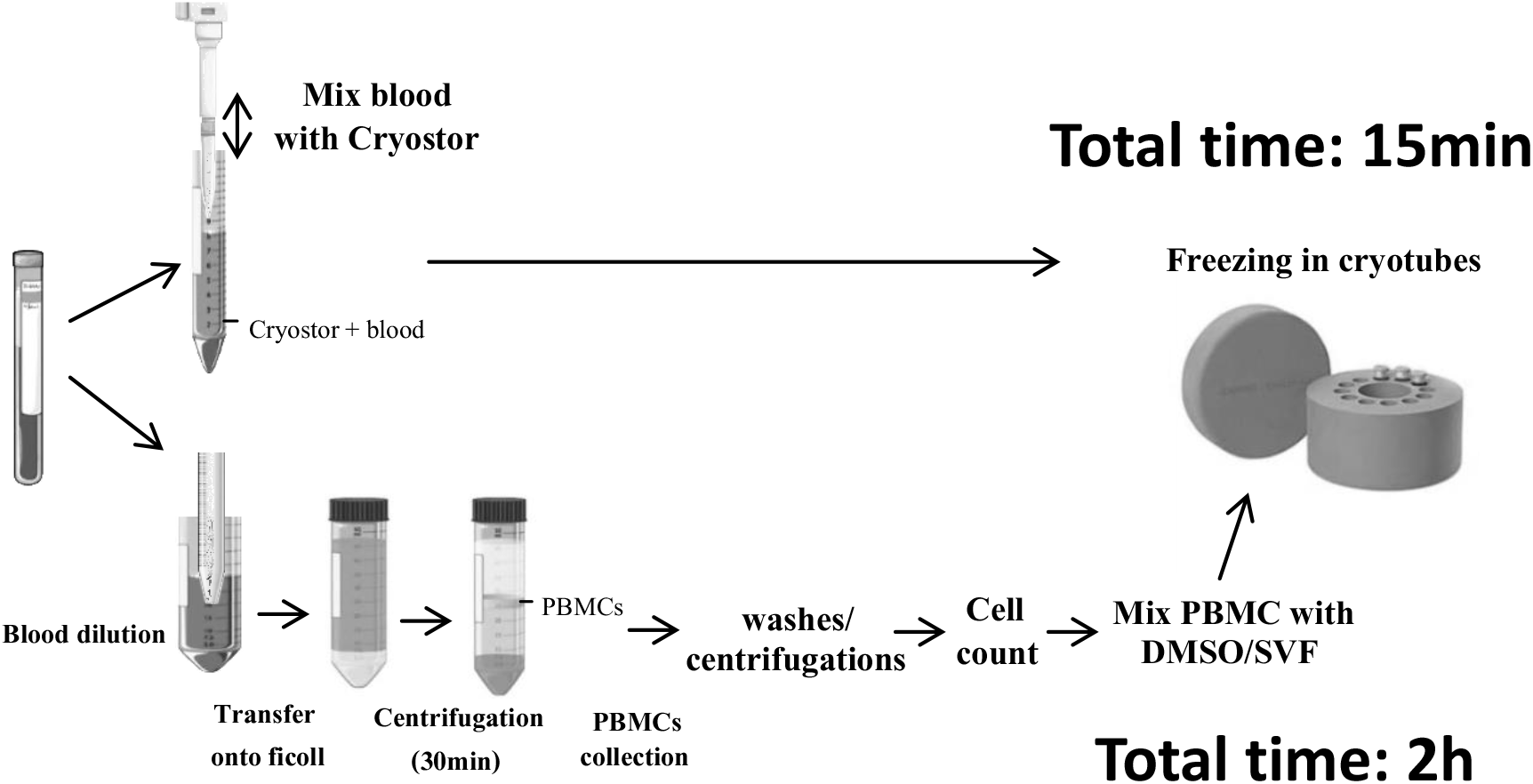
Comparison of freezing method. Whole blood was directly frozen after mixing with CryoStor^®^ CS10 (15minutes). In comparison, PBMC were isolated with a ficoll gradient, washed, counted and then frozen with 10% DMSO (2 hours).

### Good leucocytes preservation after whole blood freezing

This study established that this whole blood freezing method did not alter general morphological profile after thawing (**Figure 2A**). Only slight morphological changes were observed for whole blood leucocytes from cryopreserved vs fresh samples, which mostly accounted for a small decrease of side-scattered light (SSC) values (**Figure 2A**). In addition, a fraction of polymorphonuclear cells (PMs) exhibited reduced forward-scattered light (FSC) values in CS10 samples. FSC^low^ PMs were CD15+CD16+ live/dead^dim^ dying neutrophils (42% of total PMs) (**Figure 2B**). Dying cells have been excluded for further analyses. Accordingly, we observed a trend to increased proportions of eosinophils within total granulocytes in frozen vs fresh blood (**Figure 2C**). Basophils (Lin^-^ CD294^+^) were well preserved (0.87% frozen vs 0.71% fresh of total leucocytes; *ns*) and the viability of T cells, B cells, NK cells, monocytes was >95% in frozen blood (**Figure 2D**). As compared to fresh blood (96.4 %), total leucocyte viability was significantly reduced in frozen blood (76.3%) due to partial neutrophil loss (**Figure 2E**).

**Figure 2:**
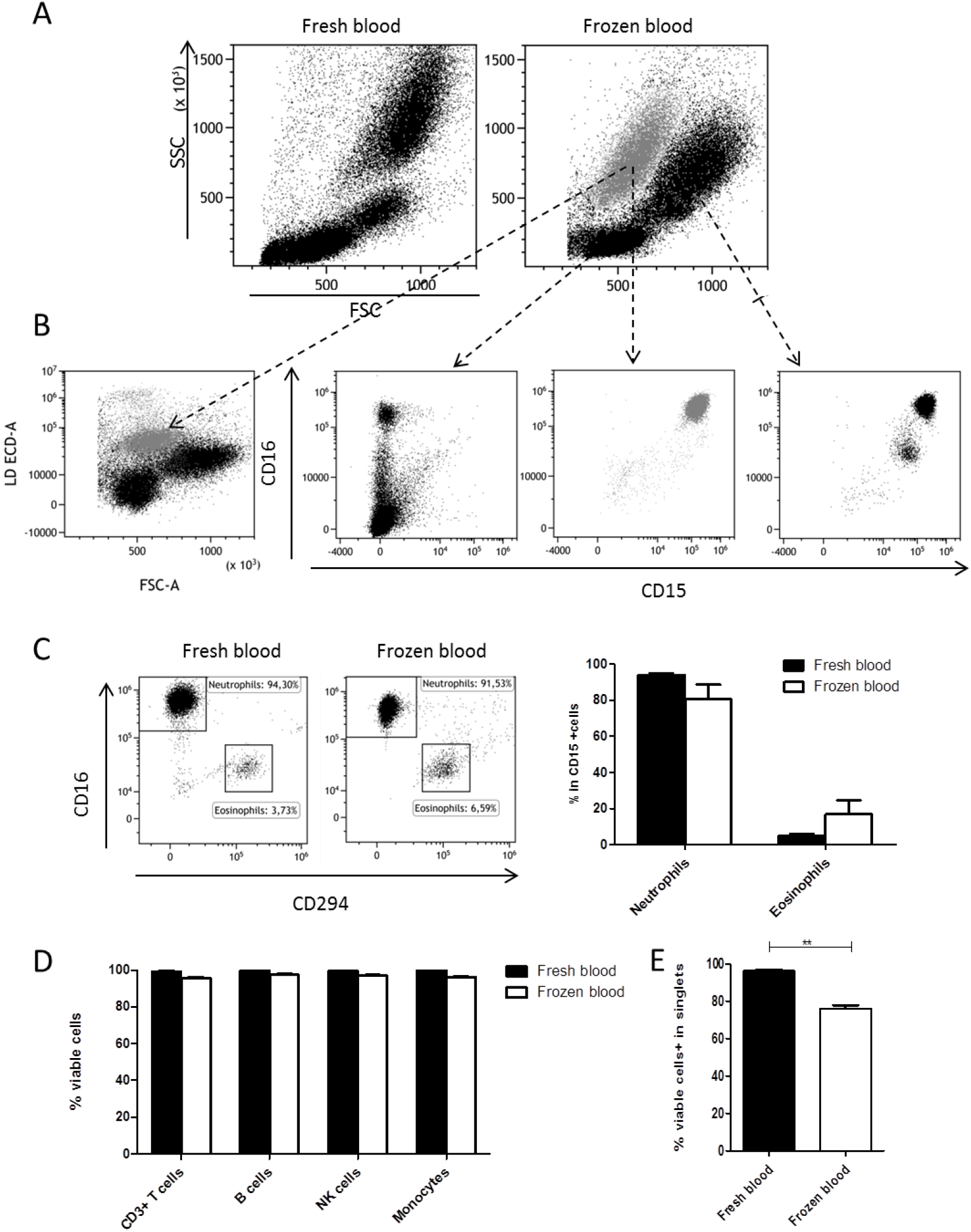
Leucocytes preservation in frozen blood. **(A)** Fresh blood and frozen blood were analyzed by flow cytometry. Representative morphologies FSC and SSC are shown. **(B)** Viability was analyzed using live/dead marker. Some granulocytic cells died (in grey), corresponded to CD15^+^CD16^+^ neutrophils. **(C)** Among living CD15^+^ polynuclear cells, neutrophils CD294^-^CD16^+^ and eosinophils CD294^+^CD16^+^ frequencies were analyzed, no statistical difference were shown between fresh and frozen blood (n=5). Results are expressed as percentage of positive cells. **(D)** Viability was >95% in T cells, B cells, NK cells and monocytes in fresh and frozen blood (n=5). **(E)** Frozen blood display 76,3% of viable cells compared to fresh blood (96,5%) (n=5)

### Immunophenotyping on cryopreserved whole blood

Similar frequencies between fresh and frozen blood were measured for CD3^+^ T cells (60.1% vs 69.5%), NK cells (mean 11.5% vs 11%) and monocytes (mean 15.1% vs 12.4%) in the lymphocyte/monocyte gate (**Figure 3A&B**). B cell frequency was decreased in frozen vs fresh blood (4.1% vs 9.7%, *p*<0.01**), nevertheless, B cell frequency in frozen blood was not significantly different from that was observed in frozen PBMC and (4.1% vs 6.6%). CD4+ CD25+ FOXP3+ Tregs were well detected in frozen blood (**Figure 4A**). A trend to slightly reduced Treg frequencies was observed in both frozen blood and PBMC as compared to fresh samples, but without reaching statistical significance, possibly due to sample number (**Figure 4B**). In addition to FOXP3, the Treg intracellular marker Helios was similarly detected across the 4 groups (**Figure 4C&D**). Total dendritic cells (DCs) (**Figure 5A**) exhibited similar frequencies in frozen vs fresh blood (1.46% vs 1.29%) as well as vs PBMC (1.46 % vs 1.54 %) (**Figure 5B**). Both CD11c+ conventional DCs (cDCs) and CD123+ plasmacytoid DCs (pDCs) were easily identified in frozen blood (**Figures 5C**), with a trend toward reduced cDC (37.5 % vs 45.9%; NS), and increased pDCs proportions in frozen vs fresh blood (58.3% vs 36.7%). All together, the expression of all surface and intracellular markers we tested in this study was well preserved in CryoStor^®^ CS10-frozen samples (Table 2).

**Figure 3:**
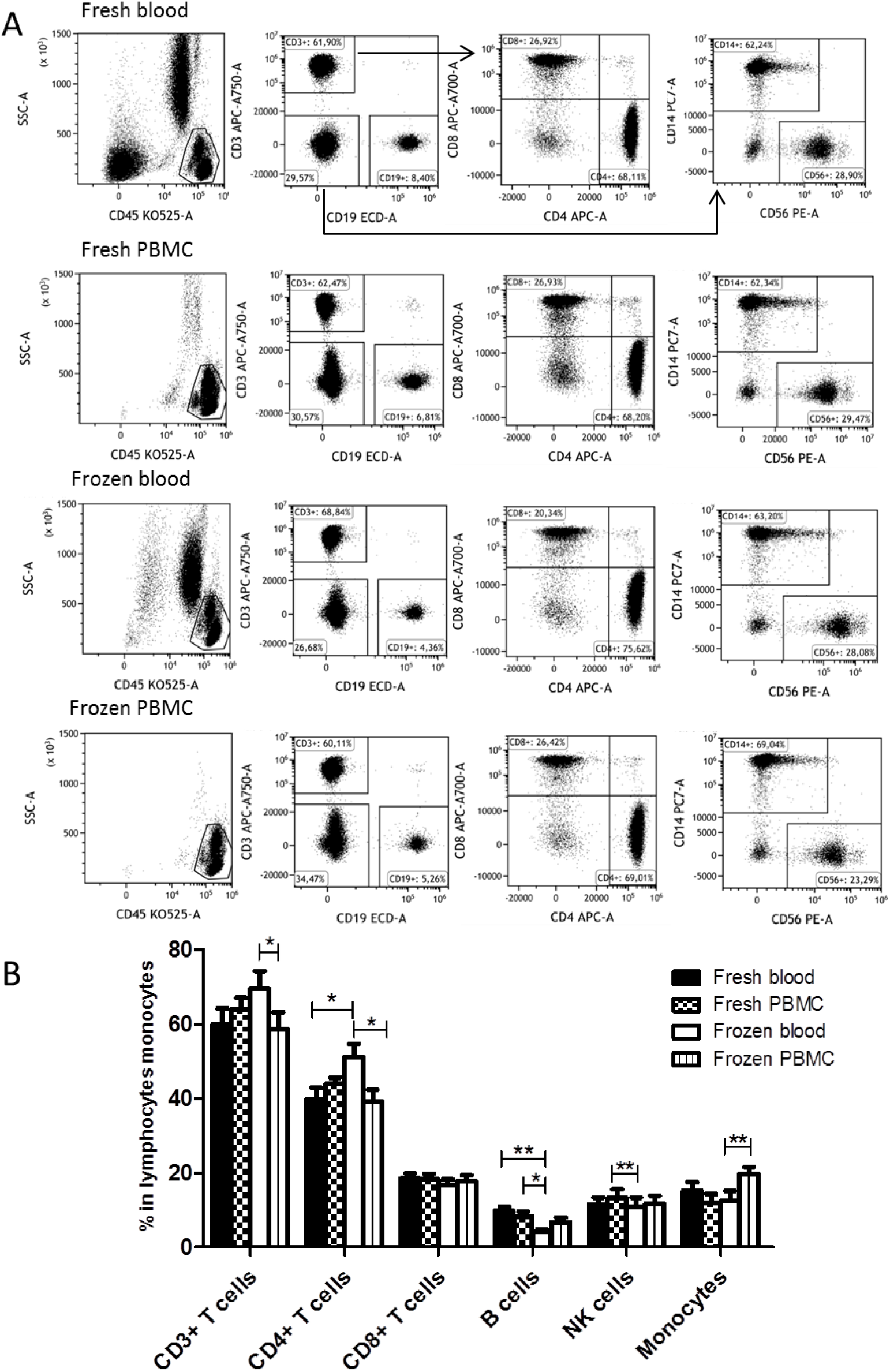
Leucocytes phenotype. **(A)** Gating strategy and representative dot plots to identify lymphoid and myeloid populations are shown. Fresh or frozen, blood or PBMC, were analyzed by flow cytometry for CD3 (T cells), CD4, CD8, CD19 (B cells), CD14 (monocytes), CD56 (NK cells) expression. **(B)** Results are expressed as percentage of positive cells (n=5).

**Figure 4:**
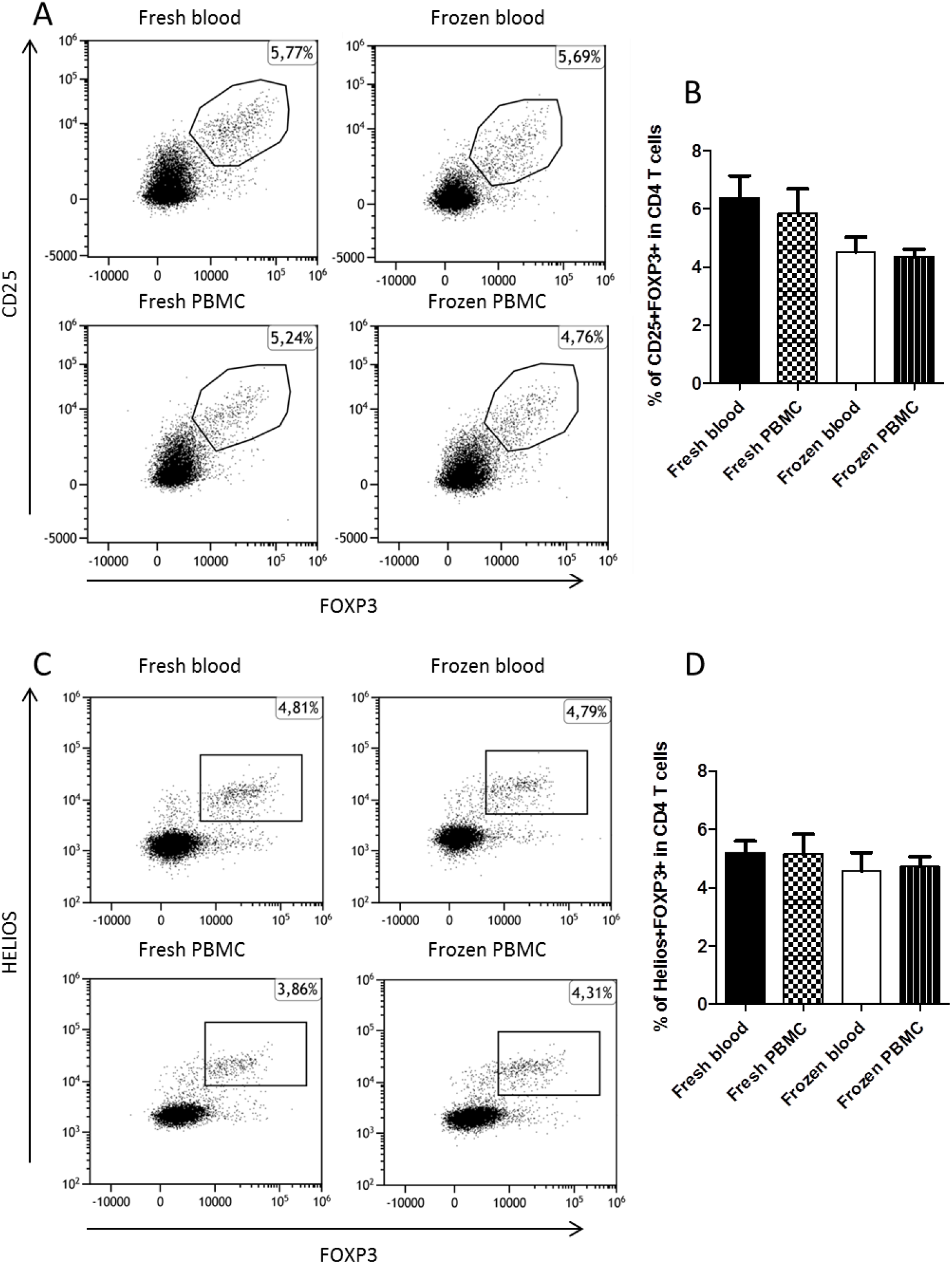
Regulatory T cells analysis. Samples were stained using Duraclone Treg tubes from Beckman Coulter. **(A and C)** Representative dot plots to identify CD25^+^FOXP3^+^ and HELIOS+FOXP3+ in CD4^+^ T cells are shown. **(B and D)** Statistical analyses for each T cells population were performed on 5 donors.

**Figure 5:**
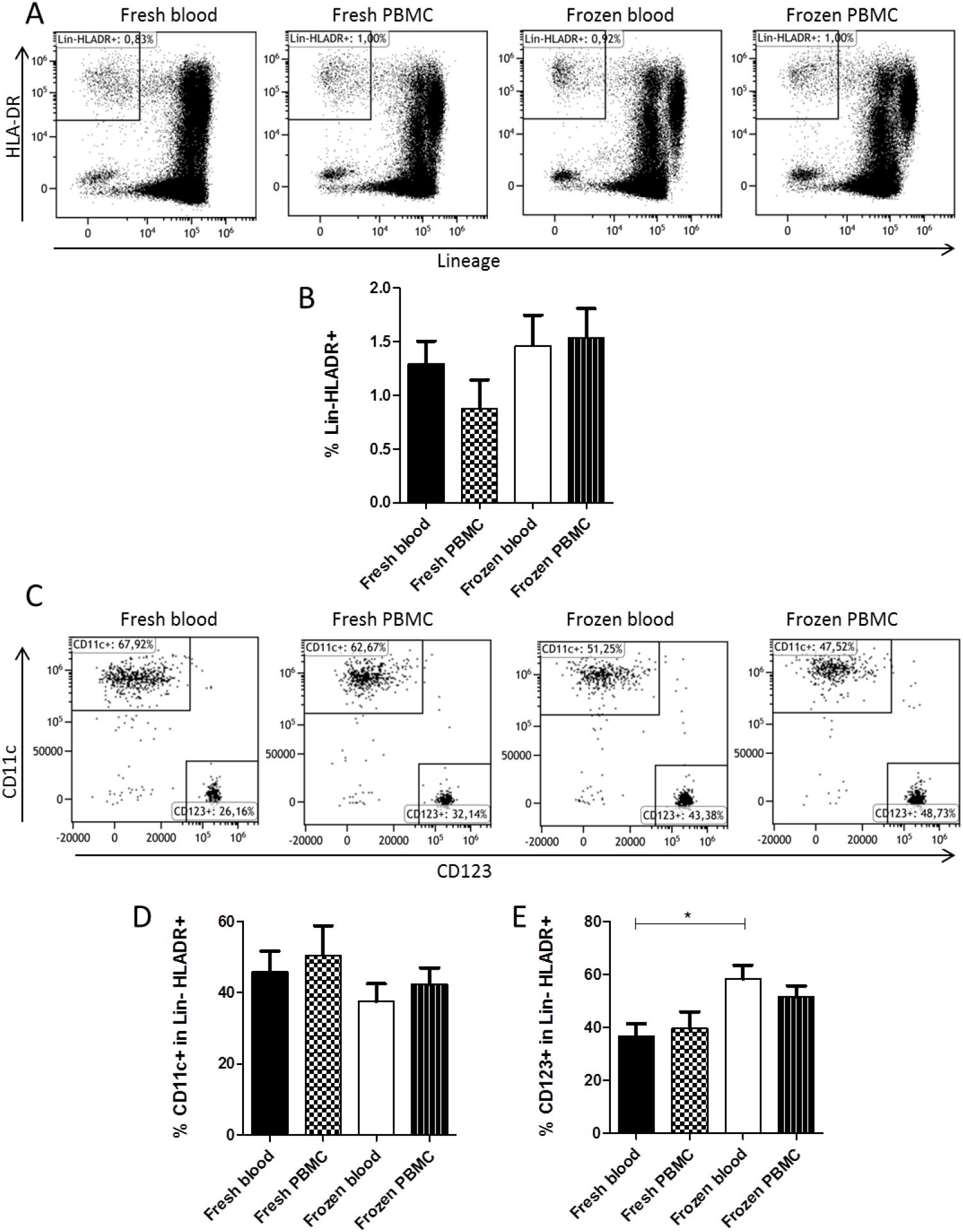
Dendritic cells analysis. Samples were stained using Duraclone dendritic cell tubes from Beckman Coulter. Representative dot plots to analyze total DCs (lineage-HLADR+) **(A) (B)** and DC subpopulation (CD11c+ or CD123+ in Lin-HLADR+) **(C) (D) (E)** are shown (n=5).

**Table 2:** Antigen expression.

| **Panel** | **Target** | **Manufacturer** | **Expression** |
| --- | --- | --- | --- |
| **Basic Panel** | CD3 | Beckman Coulter | + |
|  | CD4 | Beckman Coulter | + |
|  | CD8 | Beckman Coulter | + |
|  | CD14 | Beckman Coulter | + |
|  | CD16 | Beckman Coulter | + |
|  | CD19 | Beckman Coulter | + |
|  | CD45 | Beckman Coulter | + |
|  | CD56 | Beckman Coulter | + |
| **Dendritic cell Panel** | CD1c | Beckman Coulter | + |
|  | CD11c | Beckman Coulter | + |
|  | CD16 | Beckman Coulter | + |
|  | CD45 | Beckman Coulter | + |
|  | CD123 | Beckman Coulter | + |
|  | Clec9A | Beckman Coulter | +/- |
|  | HLA-DR | Beckman Coulter | + |
|  | Lineage | Beckman Coulter | + |
| **Granulocyte Panel** | CD11b | Beckman Coulter | + |
|  | CD15 | Beckman Coulter | + |
|  | CD16 | Beckman Coulter | + |
|  | CD33 | Beckman Coulter | + |
|  | CD45 | Beckman Coulter | + |
|  | CD62L | Beckman Coulter | + |
|  | CD274 | Beckman Coulter | + /- |
|  | CD294 | Beckman Coulter | + |
| **Regulatory T cell Panel** | CD3 | Beckman Coulter | + |
|  | CD4 | Beckman Coulter | + |
|  | CD25 | Beckman Coulter | + |
|  | CD39 | Beckman Coulter | + |
|  | CD45 | Beckman Coulter | + |
|  | CD45RA | Beckman Coulter | +/- |
|  | CD127 | BD Biosciences | + |
|  | FOXP3 | Beckman Coulter | + |
|  | Helios | Beckman Coulter | + |
| **T Helper Panel** | CD3 | Beckman Coulter | + |
|  | CD4 | Beckman Coulter | + |
|  | CD45 | Beckman Coulter | + |
| | IFNγ | Beckman Coulter | + |
|  | IL-4 | Beckman Coulter | + |
|  | IL-17a | Beckman Coulter | + |
| **Monocyte**<br>**stimulation**<br>**Panel** | CD14 | Beckman Coulter | + |
|  | CD16 | Beckman Coulter | + |
|  | CD45 | Beckman Coulter | + |
|  | IL-1b | Miltenyi Biotec | + |
|  | IL-6 | Miltenyi Biotec | + |
|  | TNFa | BD Biosciences | + |

### The function of immune cells is preserved in CS10 frozen cells

For T cell functional assays, whole blood samples were thawed and stimulated with PMA + ionomycine during 4 hours followed by intracellular staining for IL-4, IL-17A and IFNγ (**Figure 6A**). All cytokines were detected in both CD4+ and CD8+ T cells from frozen blood samples (**Figure 6B**) indicating that T cells were functional. As compared to fresh blood, there was a tendency to reduced frequencies of IFNg+ and IL-17+ cells in CD4+ and CD8+ T cells from frozen as compared to fresh blood yet without differences as compared to frozen PBMC.

**Figure 6:**
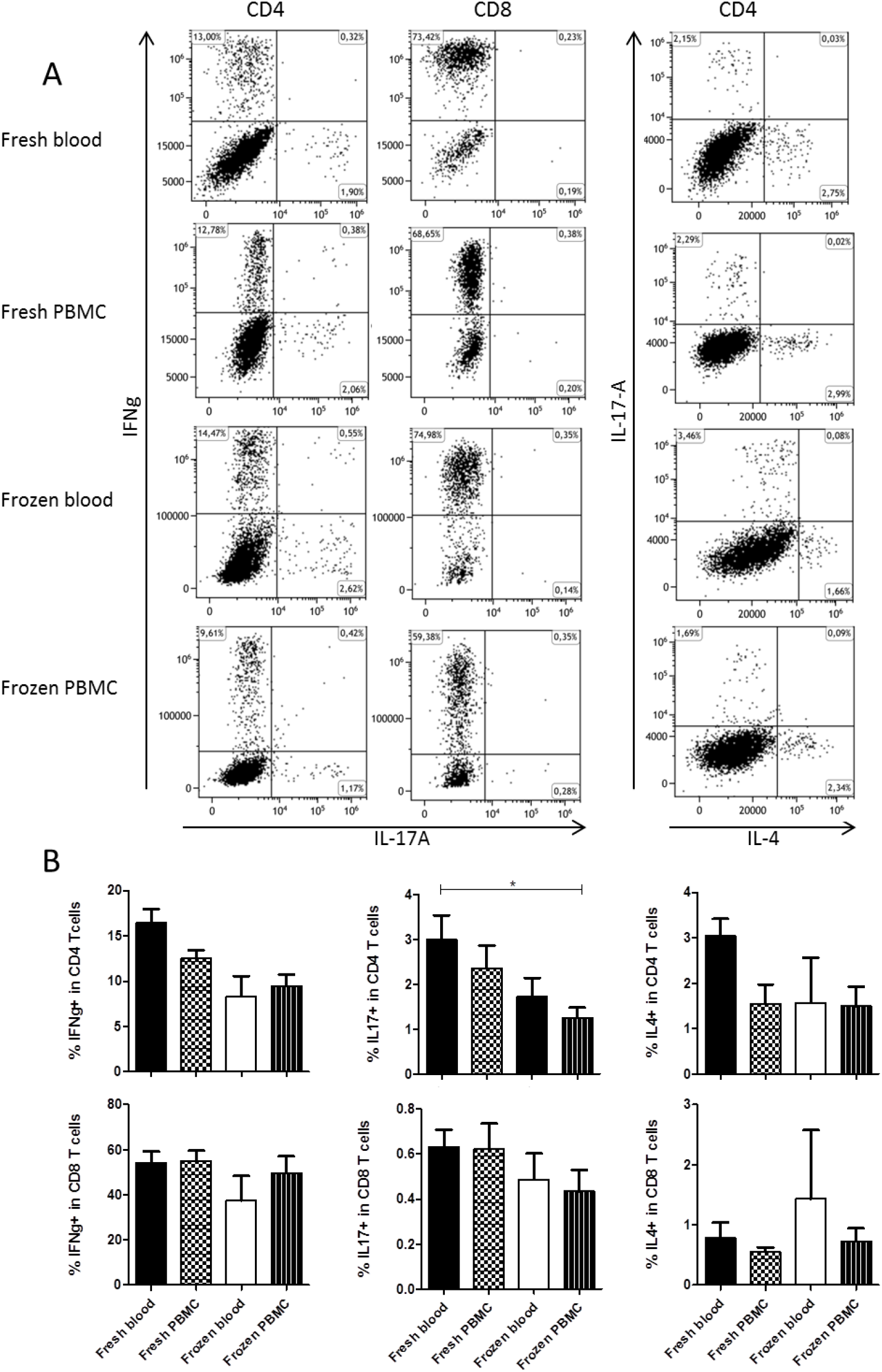
T cell function. IFNγ, IL-17A and IL-4 was assessed by flow cytometry in T cells after PMA and ionomycin stimulation of samples (n=5). **(A)** Representative examples of intracellular cytokine production are shown; **(B)** results are expressed as percentage of positive cells.

We also tested myeloid cell function by stimulating blood (fresh and frozen) with LPS during 4h followed by intracellular staining for IL-1b, IL-6 and TNFa in monocytes. As shown in figure 7, all cytokines were detected in frozen blood similarly to fresh blood (*p=ns*) indicating a good preservation of monocyte functions.

**Figure 7:**
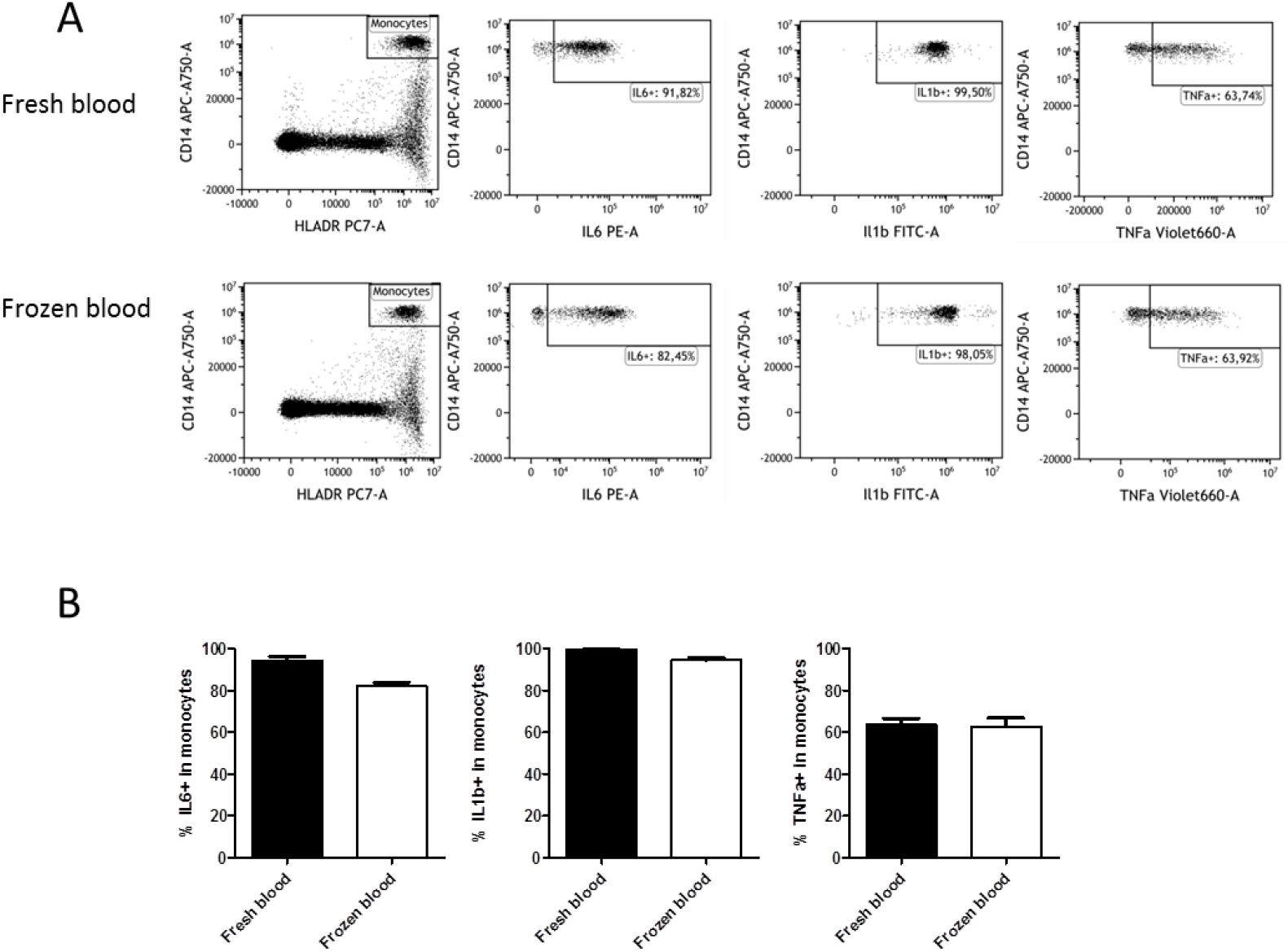
Monocyte cell function. TNFα, IL-1b and IL-6 was assessed by flow cytometry in monocytes after LPS stimulation of fresh and frozen whole blood (n=3). **(A)** Representative examples of intracellular cytokine production are shown; (B) results are expressed as percentage of positive cells.

## CONCLUSION

Here, we present a rapid and simple method for whole blood cryopreservation which allows a reliable phenotypic and functional characterization of both lymphoid and myeloid populations by flow cytometry when compared to fresh samples. Several whole blood freezing methods have been previously described. Most of the methods included fixation buffers, which resulted in poor lymphocyte viability (60-70%) (10) and with up to 50% T cell loss (11). The improved viability we observed in our study is likely explained by the optimized freezing media we used (12). Similar to two recent studies reporting new whole blood freezing methods (13,14), our method also resulted in decreased B cell abundance as compared to matched fresh blood. B cells frequency could also be affected by cryopreservation as already shown recently by Sakkestad and al (15). A major advantage of whole blood approaches is the possibility to analyze myeloid cells including PMs which are the dominant immune cell population in blood. A recent report using a similar approach as ours nevertheless indicated death rate of 60% and 32% in granulocytes and monocytes respectively (14). PMs are very sensitive to freezing and we also observed PM cell death (up to 40%) with our method, yet to a lesser extent than other published methods. Moreover, we observed very good preservation of monocytes, mDC and pDC in frozen blood.

We also validated the possibility to analyze FOXP3+ and Helios+ regulatory T cells and to perform ICS assays on T cells with similar performance as for frozen PBMC (16). Preliminary data showed that CCR7 could also be used to discriminate naïve and memory T cell subsets (data not shown). However, as chemokine receptors expression are known to be affected by freezing, the expression of other receptors of interest need to be assessed. As previously reported by Owen et al in frozen PBMC (17), cytokine production was more affected in CD4 vs CD8 T cells from cryopreserved whole blood. Our data, nevertheless, suggest the whole blood cryopreservation we describe here should be acceptable for measuring T cell responses. Interestingly we also demonstrated that myeloid cells such as monocytes and DC (data not shown) from frozen blood, were functional after freezing in CryoStor^®^ CS10 as assessed by intracellular cytokine staining. To our knowledge, this new freezing method is the first to allow functional analysis of lymphoid and myeloid cells after whole blood freezing.

Some limitations exist in this study. Analyses were performed on 5 samples from healthy donors and larger studies using samples from patients are required. Another limitation is that frozen blood samples were stored for only 3 months at −80°C before analysis. The effects of storage in liquid nitrogen and for a longer period of time need to be assessed but our preliminary results suggest a good preservation in liquid nitrogen too. Finally, B cell subsets were not assessed in this study.

The whole blood freezing method we described here is easy to use in clinical centers and does not require dedicated research staff. Another advantage is that it minimizes the volume of blood required which is especially important for pediatric populations. This technique is simple to implement and could be easily included in research design for blood immune phenotyping and functional analysis in future clinical studies.

## ACKNOWLEDGEMENTS

This work was supported by the LabEX IGO project funded by the “Investissements d’Avenir” program through the “Agence Nationale de la Recherche” (grant n° ANR11-LABX-0016-01).

